# Non-coding loci without epigenomic signals can be essential for maintaining global chromatin organization and cell viability

**DOI:** 10.1101/2020.12.13.422541

**Authors:** Bo Ding, Ying Liu, Zhiheng Liu, Lina Zheng, Ping Xu, Zhao Chen, Peiyao Wu, Ying Zhao, Qian Pan, Yu Guo, Wensheng Wei, Wei Wang

## Abstract

The majority of the non-coding regions in the human genome do not harbor any annotated element and are even not marked with any epigenomic signal or protein binding. An understudied aspect of these regions is their possible roles in stabilizing the 3D chromatin organization. To illuminate their “structural importance”, we chose to start with the non-coding regions forming many 3D contacts (referred to as hubs) and identified dozens of hubs essential for cell viability. Hi-C and single cell transcriptomic analyses showed that their deletion could significantly alter chromatin organization and impact gene expression located distal in the genome. This study revealed the 3D structural importance of non-coding loci that are not associated with any functional element, providing a new mechanistic understanding of the disease-associated genetic variations (GVs). Furthermore, our analyses also suggested a powerful approach to develop “one-drug-multiple-targets” therapeutics targeting the disease-specific non-coding regions.

## INTRODUCTION

Non-coding sequences in the human genome are known to be critical in many biological processes because they encode, such as, non-coding RNAs, enhancers and transposons that are functionally important. Despite the great progress in uncovering new roles of these non-coding elements, the dominant majority of the human genome remains unannotated. As the 3D organization of the genome is pivotal for regulating transcription and other cellular functions ^1-6^, an overlooked aspect of non-coding sequences is their “structural importance” in forming and maintaining the proper 3D chromatin structure, particularly for those that are not marked by any epigenetic signal or annotated with any functional unit.

In the analysis of protein function, residues that are not directly involved in the protein’s enzymatic activity or interaction with ligands can be important if, for example, they form the hydrophobic core to stabilize the proper conformation^7^. Similarly, non-coding genomic sequences can be critical for stabilizing the proper chromatin structure despite that they do not encode any enhancer or TF binding site. In fact, previous studies have shown that change of non-coding sequences could alter chromatin organization; for instance, deletion of some boundary sequences of topologically associating domains (TADs) ^1,2^ causes aberrant gene transcription leading to diseases ^3^. TAD boundaries can be considered as a special case but the structural importance of non-coding sequences, particularly those without associated with TAD or any functional element, has not been fully investigated.

Deleting a non-coding sequence and examining a phenotypic readout such as cell viability can directly assess its importance. High-throughput genetic screening by the CRISPR-Cas9 system has been effectively applied to analyzing non-coding RNAs^8–10^, enhancers and promoters ^11–13^. However, it is still prohibitive to delete each 5-kbp segment in the genome for a thorough screening and random selection of deletion loci is inefficient. For example, less than 3% of lncRNAs were reported to be essential ^8–10^ and this percentage is expected to be much lower for the unannotated non-coding loci. A reasonable strategy is to start from loci involved in many chromatin contacts, referred to as hubs hereinafter, because disrupting these hubs may lead to relatively profound perturbation to the chromatin organization. It is important to note that the structurally important non-coding loci may not be limited to hubs and not each hub is necessarily essential for cell viability; focusing on the hub loci in this first study simply aims to improve the identification rate for essential non-coding regions.

## RESULTS

To select the hub loci, we downloaded the 5 kb-resolution Hi-C data in 7 human cell lines (K562, GM12878, HMEC, HUVEC, IMR90, NHEK, K562 and KBM7) ^14^, and identified the significant intra-chromosome contact pairs (*p*-value cutoff of e^−20^, **Online Methods**). We next assembled all the contacts into a network, which is referred to as Fragment Contact Network (FCN) hereinafter. Each node is a 5-kb fragment and each edge represent a 3D contact.

Note that these contacts indicate the spatial closeness of the contacting loci. These contacts are not necessarily mediated by proteins or ncRNAs to form specific chromatin loops. An analogy is protein residues forming hydrophobic cores, which are located in the interior and form contacts with other residues but do not necessarily have specific residue-residue interactions mediated by such as hydrogen bonds; however, deleting these residues can disrupt the packing of the interior residues and thus distort the proper conformation required for the protein’s normal function. As an analogy, perturbing hubs may have the same impacts on the 3D genome structure by deteriorating the packing of the chromatins.

To illustrate the importance of the hubs, we first investigated their contribution to stabilizing the FCNs and association with genetic variations in cancer. Then we identified hubs essential for cell viability using a CRISPR screening. Finally, we illustrated the impact on chromatin structure and gene expression of hub deletion using Hi-C and single cell RNA-seq.

### FCN networks are resistant to random attacks but vulnerable to targeted attacks

Due to the sparse inter-chromosome contacts that can be called from Hi-C, we focused on the intra-chromosome contacts and constructed FCN for each chromosome in each cell line, resulting in total 161 (=23*7) FCNs for all chromosomes in the 7 cell lines. We found that the degree distribution of FCN follows a power-law (**Fig. 1a**), indicating that FCNs are scale-free networks. FCNs are in fact resistant to random attack but vulnerable to targeted attack to high degree nodes, as all scale-free networks. The 161 FCNs have similar network parameters, such as effective diameters, which is the path length such that 90 percent of node pairs are at a smaller or equal distance apart (see FCN Network Analysis Results in **Supplementary Materials**). The most significant outlier was FCN of chr9 in the leukemia cancer cell line K562, which had a significantly larger effective diameter than the rest (**Fig. 1b** and **Supplementary Fig. 1a**). We found that computationally removing high-degree nodes in chr9 of GM12878 led to a similar degree distribution of chr9 in K562 (**Fig. 1c**). This analysis suggested genetic variations in K562 likely target the high-degree nodes in chr9 and thus alter the network properties. We also confirmed that the high degree nodes (hubs) are crucial in stabilizing the contacts between their connecting nodes in the network (defined as “neighbors”, hereinafter) (**Supplementary Fig. 1c-f**).

**Figure 1.**
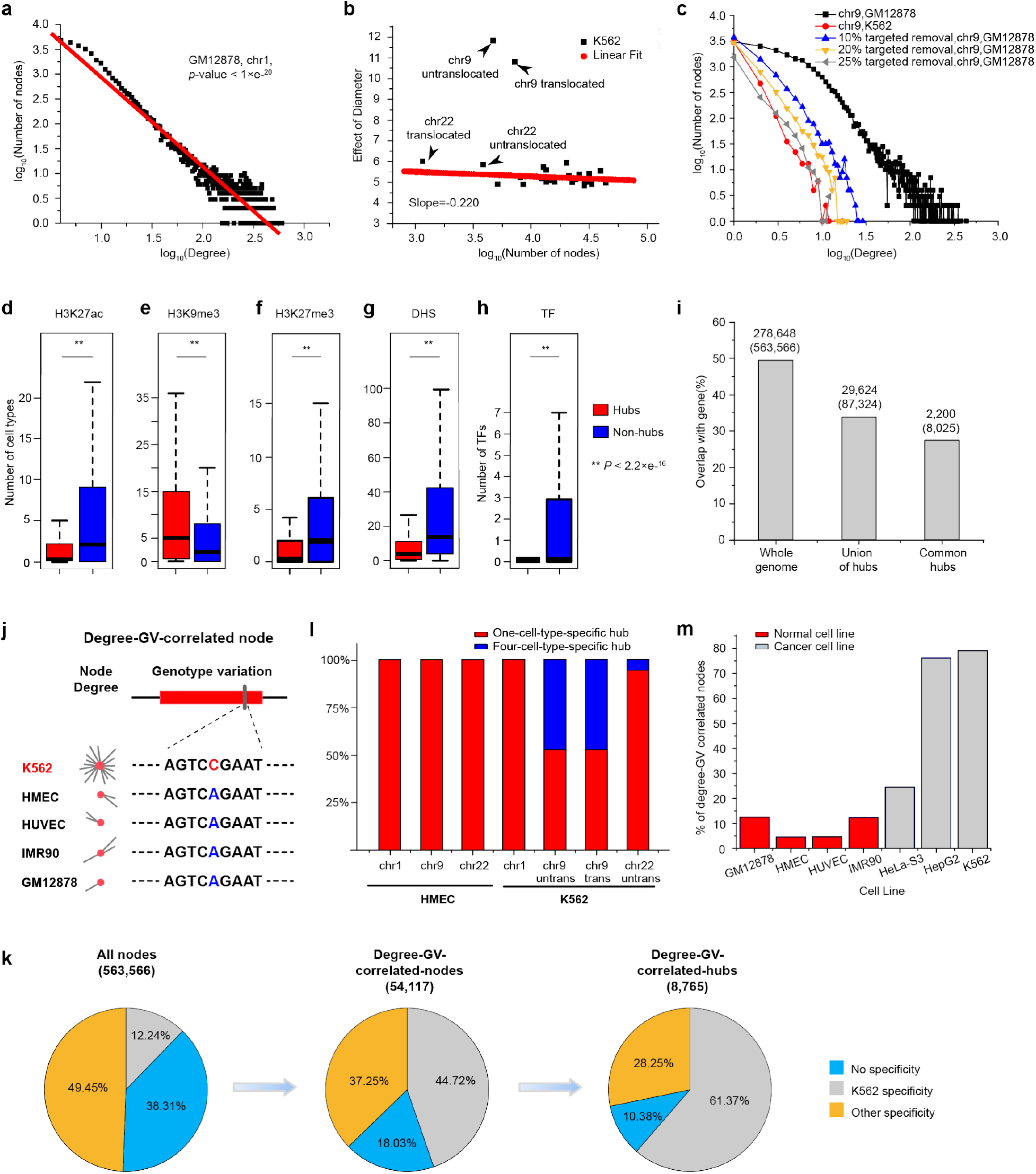
Characterization of the FCNs and hub nodes. (**a**) The degree distribution of FCN composed of Hi-C contacts with *p*-value < e^−20^. (**b**) The effective diameter of FCN remains largely unchanged with the increase of network size. Chr9 in K562 is the only outlier of all chromosomes with a large effective diameter, while chr22 is similar to others. (**c**) The degree distribution of chr9 in normal cell lines after targeted removal of high-degree nodes is similar to that of K562 chr9. Chr9 of GM12878 is shown as an example with its top 10%, 20% and 25% of highest connected nodes removed. (**d-h**) Comparison of hub and non-hub in terms of H3K27ac (**d**), H3K9me3 (**e**), H3K27me3 (**f**), DNaseI hypersensitivity sites (DHS) (**g**) and TF ChIP-seq peaks (**h**) in 121 cell lines/primary cells/tissues characterized by the NIH Roadmap Epigenetics Project. Other epigenetic marks are shown in **Supplementary Fig. 1**. (**i**) The percentages of gene coding regions in the whole genome, union of hubs (hubs appeared in at least one cell type) and common hubs (hubs appeared in all cell lines). Numbers above the histogram are the number of nodes overlapped with gene coding regions and the total number of nodes in that category (in parenthesis). (**j**) Definition of degree-GV-correlated nodes. In this example, the node has high degree in K562 and low degree in other cell lines, which is correlated with the GV profile with a SNP in K562 but none in other cell lines. (**k**) The distribution of all cell-line specificities in All-nodes, Degree-GV-correlated-nodes and Degree-GV-correlated-hubs. Cell line specificity suggests that the contact degree was specifically high or low in any particular cell line (**Online Methods** and **Fig. 1j**). (**I**) The distribution of one-cell-specific hubs and four-cell-specific hubs in chromosomes and cell lines. Chr9 in K562 is significantly enriched with hubs commonly found in other 4 cell types but not in K562 (four-cell-specific hubs). All the data are shown in (**Supplementary Table 6**). (**m**) The percentage of Degree-GV correlated nodes in normal cell lines and cancer cell lines.

Compared to non-hubs, hubs are often marked by H3K9me3 but lack peaks of all the other histone marks. Consistently, we observed less TF binding, less open chromatin (**Fig. 1d-h** and **Supplementary Fig. 1g-i**), and fewer annotated genes in hubs than in non-hub regions (**Fig. 1i**). These observations suggested that hubs are similar to the hydrophobic cores in proteins, both densely packed in the interior of the 3D structure. Therefore, perturbation to hubs by such as genetic variations and deletions could disrupt the chromatin packing to affect the surrounding 3D organization of the chromatin, and propagate through the genome and lead to observable phenotypes such as disease formation and cell death. Next, we examined whether the hubs found in the normal cells have significantly different 3D contacts in cancers and whether such change is associated with genetic variations (GVs)(referred to single nucleotide variation (SNV) hereinafter). Then, we investigated whether and how deleting hubs can cause cell death.

### Cancer related mutations alter 3D contacts of hubs

As K562 is a cancer cell line, we investigated whether K562-specific genetic variations (GVs) are related to the changes of spatial contacts. After calculating Z-score for each node’s degree so that it is comparable across cell lines, we checked its specificity, i.e. whether the contact degree was specifically high in any particular cell line (**Online Methods** and **Fig. 1j**). When considering all the nodes, we did not observe any specificity bias towards K562: in the total 563,566 nodes of the whole genome, 38.3% showed no specificity, 12.2% K562-specific, and the largest of other specificities was 13.4% (**Fig. 1k** and **Supplementary Table 4**).

Next, we calculated the Pearson correlation coefficient between a node’s degree and the GV occurrence in the node across cell lines. We found that K562-specific GVs were associated with the degree changes in K562: among all the 54,117 nodes with degree-GV *Pearson* correlation coefficients > 0.9 (referred to as degree-GV-correlated nodes, **Fig. 1j**), 24,229 (44.7%) were K562-specific; as a comparison, the largest percentage for another cell-type (HMEC) specificity was only 10.6% (5,743 nodes) (**Fig. 1k** and **Supplementary Table 4**). This bias towards the only diploid cancer cell K562 among the seven was even more obvious for hubs: for all the hubs identified in at least one of the 7 cell lines, there were 8,765 degree-GV-correlated hubs, among which 5,379 (61.4%) were K562-specific compared to the largest percentage of 824 (9.4%) specific to another cell type (HMEC) (**Fig. 1k** and **Supplementary Table 4**). Taken together, these analyses suggested that K562-specific GVs tend to significantly change the contact degrees, particularly on hubs, which is consistent with the observation that the FCN is vulnerable to targeted genetic variations in hubs.

Genetic variations can either disrupt hubs in the normal cells or form new disease-specific hubs in cancer cells. We thus analyzed hub formation and disruption separately, and found strong correlation between GV and contact degree change in K562 for both scenarios (see FCN Network Analysis Results in **Supplementary Materials**). In particular, the percentages of hub disruption in chr9 of K562 (i.e. hubs found in the other 4 cell types but not in K562) were 47.56% and 47.50%, respectively, without and with considering translocation between its chr9 and chr22 (only the untranslocated part of chr9 used for calculation). This was significantly higher than all other chromosomes in each cell line, whose range is between 0% and 18.6% (**Fig. 1l**). Our analyses clearly showed that GVs in K562 severely disrupted the hubs shared by other cell lines in chr9.

To confirm the generality of this observation, we extended our analysis to 4 normal (GM12878, HMEC, HUVEC and IMR90) and 3 cancer cells (HepG2, HeLa-S3 and K562) that had both 20-kb resolution Hi-C and GV data. We also found strong correlation between contact degree and GV in cancers (**Fig. 1m**), suggesting that the cancer-specific GVs tend to significantly alter the 3D contacts of hubs.

### Targeted deletion on hubs can significantly affect cell viability

The above analyses indicated that hubs are not necessarily directly involved in functional activities such as hosting enhancers, but they can be crucial for stabilizing the chromatin structure and thus are functionally important. To further test this hypothesis, we selected 960 hub regions (each 5-kbp in length) to examine their impact on cell viability in a high-throughput deletion screen (**Supplementary Table 8**). These hubs are those likely to stabilize the contacts between neighbors (see FCN Network Analysis Results in **Supplementary Materials**), including 683 hubs present in all cell lines and 277 hubs specific for K562 cells.

For the screening, we constructed a paired-gRNA (pgRNA) library ^8^ targeting the selected hubs mediated by CRISPR-Cas9 system. Using lentiviral transduction at a low MOI (multiplicity of infection) of < 0.3, we respectively transfected the pgRNA library containing a total of 17,476 pgRNAs into K562 cells stably expressing Cas9 protein. This library also included 473 pgRNAs targeting essential ribosomal genes as positive controls, 100 pgRNAs targeting *AAVS1* locus and 100 non-targeting pgRNAs as negative controls (**Supplementary Table 9**). The library cells were cultured for continuous 30 days post transduction. We sequenced cells at Day-0 (controls) and Day-30 to determine the abundance of barcode-gRNA regions, which represent the corresponding pgRNAs (**Fig. 2a**).

**Figure 2.**
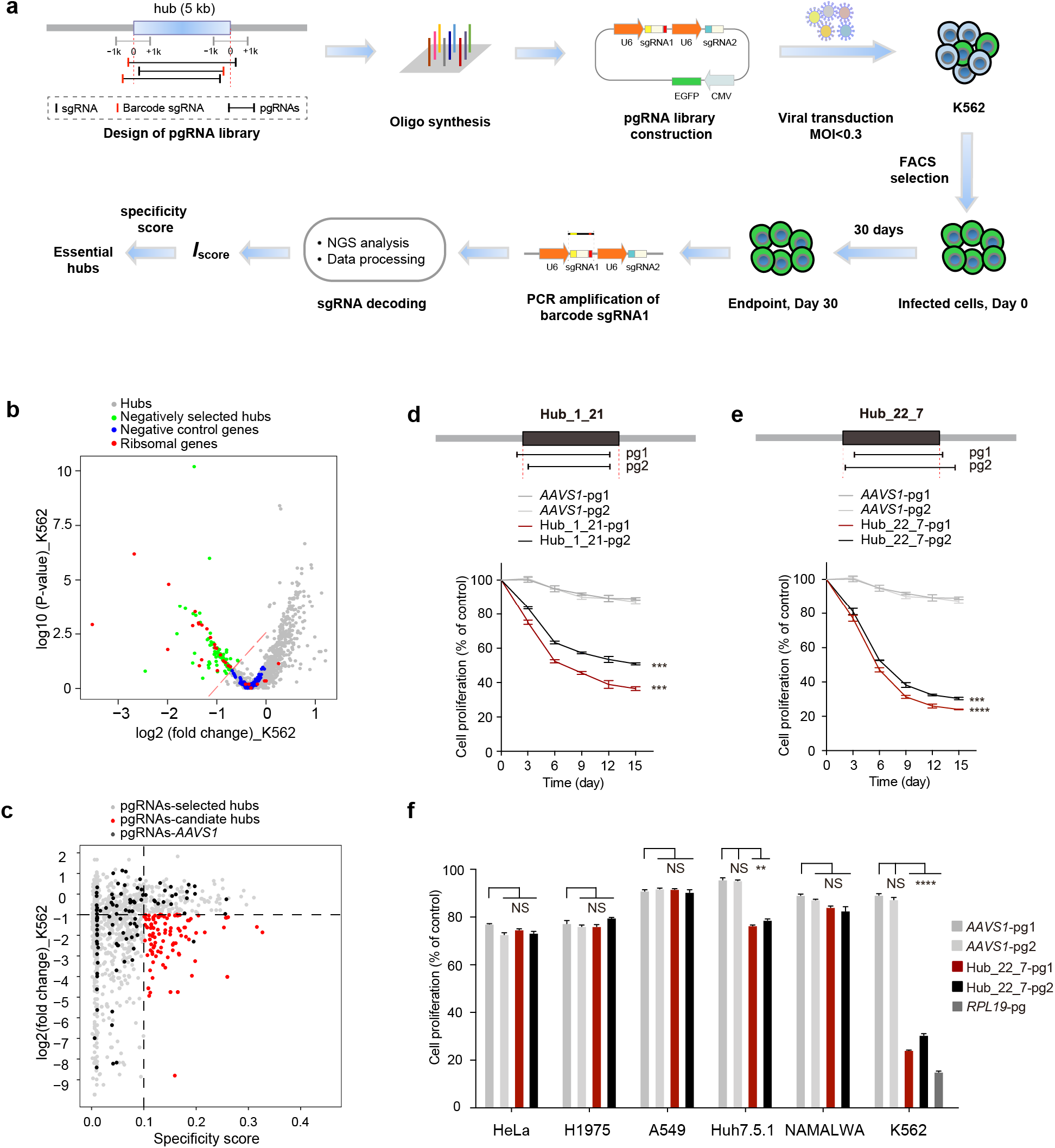
Identification of essential hubs for cell growth and proliferation in K562 cell line through pgRNA-mediated fragment deletion. (**a**) The Schematic of the pgRNA library design, cloning and functional screening of selected hub loci. (**b**) Volcano plot of the fold change and *p-value* of hubs in the K562 cell line. Negative control genes were generated by randomly sampling 20 *AAVS1* - targeting pgRNAs with replacement per gene, and essential genes represent ribosomal genes serving as positive control in the screening. Dotted red line represents *Iscore* = −1. (**c**) Selection of candidate essential hubs by pgRNA fold change and specificity score. These essential hits were selected under the threshold of specificity score > 0.1, log_2_ (fold change) < −1. (**d-e**) Validation of top-ranked essential hubs in K562 cell by cell proliferation assay. *AAVS1*-pg1 and *AAVS1*-pg2 are pgRNAs targeting *AAVS1* locus as negative controls. Cells were individually infected with lentivirus of pgRNA vector containing an EGFP marker. The percentages of EGFP-expressing cells indicating the fraction of pgRNA-containing cells were quantified every 3 days by FACS. Cell proliferation of each sample was measured by normalizing the percentage of EGFP^+^ cells at each time point with that at 3 days post infection (labeled as Day 0). Asterisk (*) represents *p-value* compared with pgRNAs targeting *AAVS1*-pg1 at Day 15, calculated by two-tailed Student’s *t*-test and adjusted for multiple comparisons by Benjamini–Hochberg procedure. Data are presented as the mean ± s.d. (n = 3 biologically independent samples). \**p* < 0.05; \*\**p* < 0.01; *** *p* < 0.001; **** *p* < 0.0001; NS, not significant. The pgRNAs for individual validation of each hub locus are listed in **Supplementary Table 12**. (**f**) Validation of hub_22_7 in multiple cancer cell lines including A549, H1975, HeLa, Huh7.5.1 and NAMALWA cell lines. Asterisk (*) represents *p-value* compared with pgRNAs targeting *AAVS1*-pg1 at Day 15, calculated by twotailed Student’s *t*-test and adjusted by Bonferroni correction accounting for multiple testings. * *p* < 0.05; \*\**p* < 0.01; *** *p* < 0,001; \*\*\*\**p* < 0.0001; NS, not significant.

Distributions of pgRNA reads in two biological replicates are highly correlated (**Supplementary Fig. 3a-c**). In Day-30 cell population, compared with non-targeting pgRNAs or those targeting *AAVS1,* we identified hub regions with significant depletion in their targeting pgRNAs, consistent with positive controls that target ribosomal essential genes. The fold changes of all pgRNAs targeting each hub were calculated and their *P* values were computed by comparing with the *AAVS1*-targeting pgRNAs using Mann-Whitney U test. *AAVS1*-targeting pgRNAs were randomly sampled to generate a distribution of negative controls, which was used to compute the hubs’ *P* values. Combining the mean fold change and corrected *P* values, a screen score was computed for each hub, and the hubs whose screen scores were less than or equal to −1 were considered as essential hits (**Online Methods** and **Supplementary Table 10, 11**). Overall, 77 hubs were thus selected in K562, whose depletion led to cell death or growth inhibition (**Fig. 2b**).

It has been reported that multiple cleavages in genomic loci generated by Cas9 activity could lead to cellular toxicity and thus affect the growth screen measurements ^15–18^. To minimize the potential off-target effects, we calculated the GuideScan specificity score ^19^ for each sgRNA of every pgRNA, which focused on assessing the specificities of sgRNAs with 2 or 3 mismatches to off-target loci that are commonly used in library screens, and generated a specificity score for each pgRNA. We found that pgRNA targeting *AAVS1* with specificity score ≤0.1 could lead to significant dropout effect in K562 (**Supplementary Fig. 3d**). To further assure the targeting specificity, we only selected the targeting pgRNAs with specificity score > 0.1 and log_2_ (fold change) < −1 for subsequent analysis (**Fig. 2c**). Furthermore, hub loci with copy number amplification were also filtered to minimize the effect due to multiple cleavages by certain pgRNAs^20^. Using these stringent criteria, we identified 35 essential hubs in K562 (**Fig. 2c**).

We then chose 7 candidate hubs for individual validation in K562. For each hub, two or three pgRNAs with high specificity scores were selected (**Online Methods** and **Supplementary Table 12**). All but two identified hubs were validated to severely affect cell growth and proliferation in K562 (**Fig. 2d-e** and **Supplementary Fig. 4**), indicating their functional roles in cell fitness. To further explore the cell type specificity of the essential hubs, we selected hub_22_7 (chr22: 17,325,000-17,330,000, hg19) that showed the most significant growth defect in K562, and performed the same cell proliferation assay in 5 other cancer cell lines. Interestedly, compared with negative controls targeting *AAVS1* locus, targeted deletion of hub_22_7 did not lead to significant cell death or cell growth inhibition in 4 tested cell lines including HeLa (cervical cancer cells), H1975 (non-small cell lung cancer cells), A549 (non-small cell lung cancer cells) and NAMALWA (Burkitt’s lymphoma) (**Fig. 2f** and **Supplementary Fig. 5a**). In a liver cancer cell line Huh7.5.1, deletion of hub_22_7 showed weak effect in cell fitness compared with deleting the essential gene *RPL19* serving as the positive control (**Fig. 2f** and **Supplementary Fig. 5a**). In all, hub_22_7 locus only exhibited a remarkable essential role in the K562 cell line.

### Cell death caused by hub deletion is not resulted from disruption of functional elements or off-target effect

To illuminate the mechanism of cell death induced by hub deletion, we first examined the functional annotation and epigenetic modifications in these regions. None of the essential hubs overlap with gene coding regions, non-coding RNA regions or TAD boundaries. Importantly, 77.1% (27 out of 35 including 3 out of 5 individually validated ones) of the essential hubs do not overlap with any histone modification or TF ChIP-seq peak (**Fig. 3a**, an example of hub_22_7 in **Fig. 3b** and **Supplementary Fig. 6**), indicating that the essentiality of these hubs is not resulted from the genes or regulatory elements they harbor.

**Figure 3.**
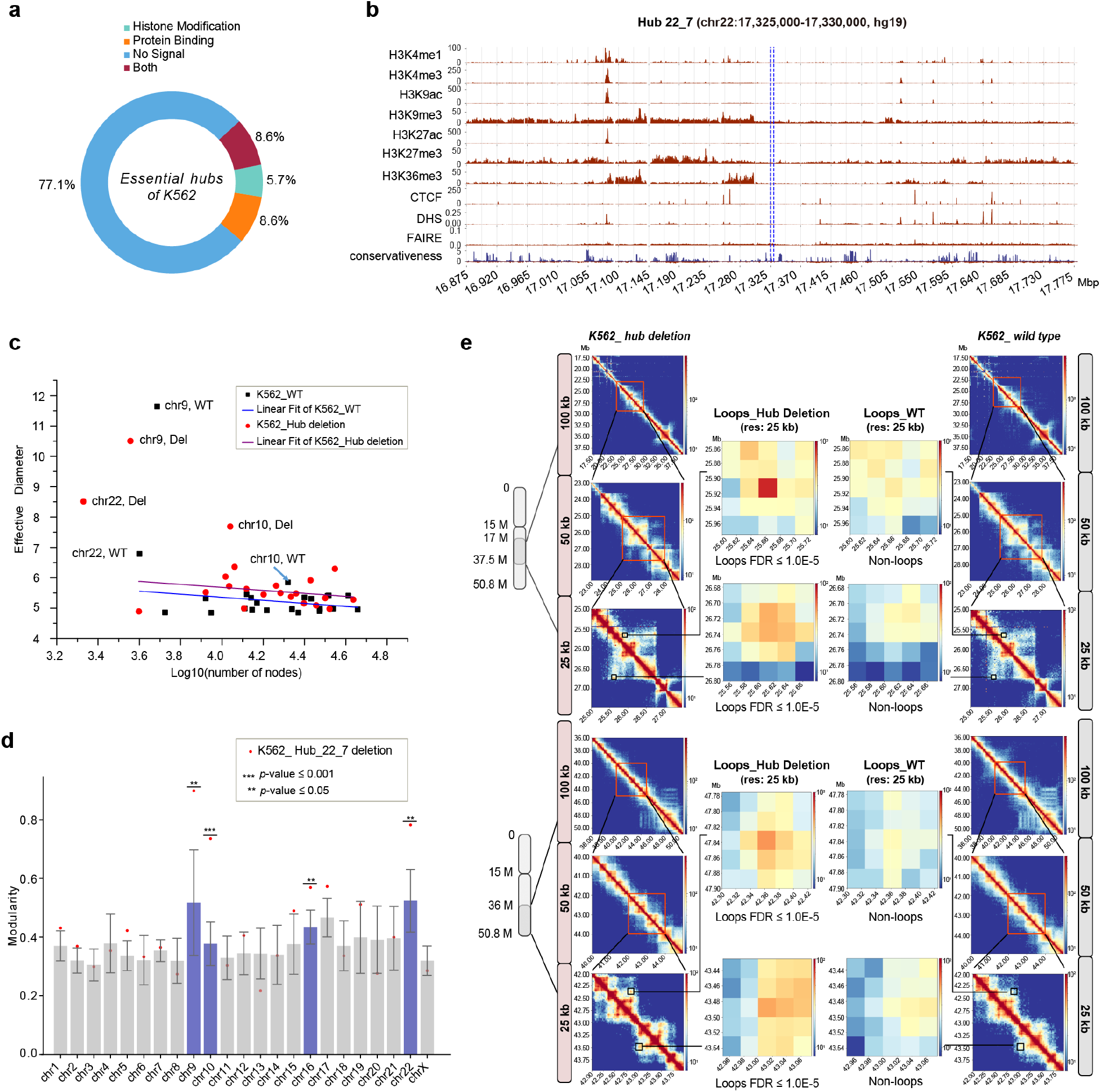
Characterization of the deleted hub and the impact of its deletion on the global chromatin structure. (**a**) Overlap of essential hubs in K562 with the peaks of 10 histone marks (H3K27ac, H3K4me1, H3K4me3, H3K27me3, H3K9me3, H3K36me3, H3K4me2, H3K79me2, H3K9ac, H3K20me1) and 151 TFs. (**b**) Histone marks, CTCF binding, open chromatin and conservation score (100 vertebrates Basewise Conservation by PhyloP) in chr22: 17,325,000-17,330,000 (see **Supplementary Fig**. **6** for all signals). (**c**) Effective diameters vs log10(number of nodes) the wild type and hub_22_7-deleted K562 cells. (**d**) Modularity scores in the 7 wild type cell lines for 23 chromosomes. Red dots: hub_22_7 deletion. (**e**) Hi-C contacts (100-kb, 50-kb and 25-kb resolution) and examples of chromatin loops (called at 25-kb resolution) in hub-deleted and wild type K562 cells.

We next evaluated the essential hub_22_7 to rule out the possibility of cell death caused by off-target cleavage. Using the validated pgRNA hub_22_7-pg2 with high specificity (**Supplementary Table 12**), we firstly measured its deletion efficiency by real-time qPCR (**Supplementary Fig. 5b**) on each time point post pgRNA transduction, then performed whole-genome sequencing (WGS) to evaluate its potential off-target effect on the day showing the highest deletion efficiency (**Online Methods**). We identified > 3.7 million SNVs and > 890,000 indels compared to the hg19 reference genome (**Supplementary Table 13**). The fact that we could successfully identify 87.4% germline mutations found in the published wild type K562 cells (ENCODE database with the accession code ENCFF313MGL, ENCFF004THU, ENCFF506TKC and ENCFF066GQD) suggests a reliable library quality. We manually checked the indels on 455 potential off-target loci and 2 on-target loci identified by Cas-OFFinder^21^ using loose criteria (bulge = 0, mismatch ≤4; bulge ≤ 2, mismatch ≤ 2) to avoid missing any possible off-target site (**Online Methods**). Significant indels were only found in 2 on-target loci and not found in any off-target loci, indicating no off-target cleavage. These analyses confirmed that cell death caused by hub deletion was not resulted from off-target effect.

### Deletion of essential hubs alters the global chromatin structure

We next performed Hi-C analysis to examine the chromatin structure changes in the hub_22_7-pg2-infected (hub_22_7-deleted) K562 cells. To characterize the global impact of hub deletion, we first constructed FCNs in the hub_22_7 deleted cells using the same criteria as in the wild type and analyzed the change of the network properties including effective diameter and modularity, which is the difference between the fraction of edges observed within a group of nodes and the expected value in random network.

By analyzing the effective diameters of the FCNs before and after the hub deletion, we found chr9, chr10 and chr22 had significant changes (*p*-value < 0.05, **Online Methods** and **Fig. 3c**): chr22 and chr10 increased while chr9 decreased upon hub deletion. The hub-deleted cells also showed significant changes of modularity scores of chr9, ch10, chr16 and chr22 (*p*-value < 0.05, **Online Methods** and **Fig. 3d**). While the change of the hub’s residing chr22 and the chr22-translocated chr9 in K562 is not unexpected, the surprising impact on chr10 and chr16 illuminates the importance of the understudied interactions between chromosomes (**Supplementary Fig. 7**).

The increased diameter and modularity in chr22 suggest that hub deletion reduces long-range chromatin contacts and enhances modularization of the FCN, consistent with the overall Hi-C contact difference between the wildtype and hub deleted cells (**Fig. 3e**). We did find newly formed chromatin loops in hub deleted cells (examples in **Fig. 3e**). Taken together, deletion of a hub has a global impact on chromatin structure that can propagate to other chromosomes.

### Deletion of essential hubs upregulates apoptotic genes

Next, we set out to identify genes whose expressions were significantly affected by hub deletion. Cells transduced with pgRNAs have various rates towards cell death and the cell population is thus heterogeneous. Therefore, we used single cell analysis to define the different cell states in the population. We performed Drop-seq analysis ^22^ on the hub_22_7-pg2 infected K562 cells and collected scRNA-seq data for 393 cells passing the quality control. The bulk RNA-seq data of the wild type and *AAVS1* deletion in K562 were included as controls. All the RNA-seq data were normalized using counts per million (CPM) and calculated the scaled z-score for each gene in each individual cell or bulk sample by fitting a binomial distribution Binomial *(N, Pi)* (**Online Methods**). The scaled z-score matrix of single cell and bulk RNA-seq data were used for the following analysis.

We performed trajectory branching and pseudotime analysis using Monocle^23,24^. Because hub deletion caused significant cell death, we selected and analyzed 93 apoptosis genes documented in the KEGG database ^25^. The single cells together with bulk samples of the wild type and *AAVS1* deletion were grouped into 5 cell states (**Fig. 4a**). Both *AAVS1* deletion and wild type were assigned to state 1, indicating that *AAVS1* deletion is a valid control. Single cells in state 1 resemble the wild type cells at the low value of pseudotime, which is understandable because hub deletions were not synchronized in all cells. State 4 and 5 have the highest pseudotime values and thus are most distinct from the wild type. Overall, the apoptosis genes showed increasing expression levels from state 2 to state 5 (see examples in **Fig. 4b**). Because states 2, 4 and 5 are the terminal nodes in the trajectory tree that represent local minimum or maximum points, we clustered the apoptosis genes according to their expression profiles in these three states. Each of the gene clusters presented unique patterns towards cell apoptosis (**Fig. 4c**). This single cell RNA-seq analysis depicted the transcriptomic progression towards cell death upon hub deletion.

**Figure 4.**
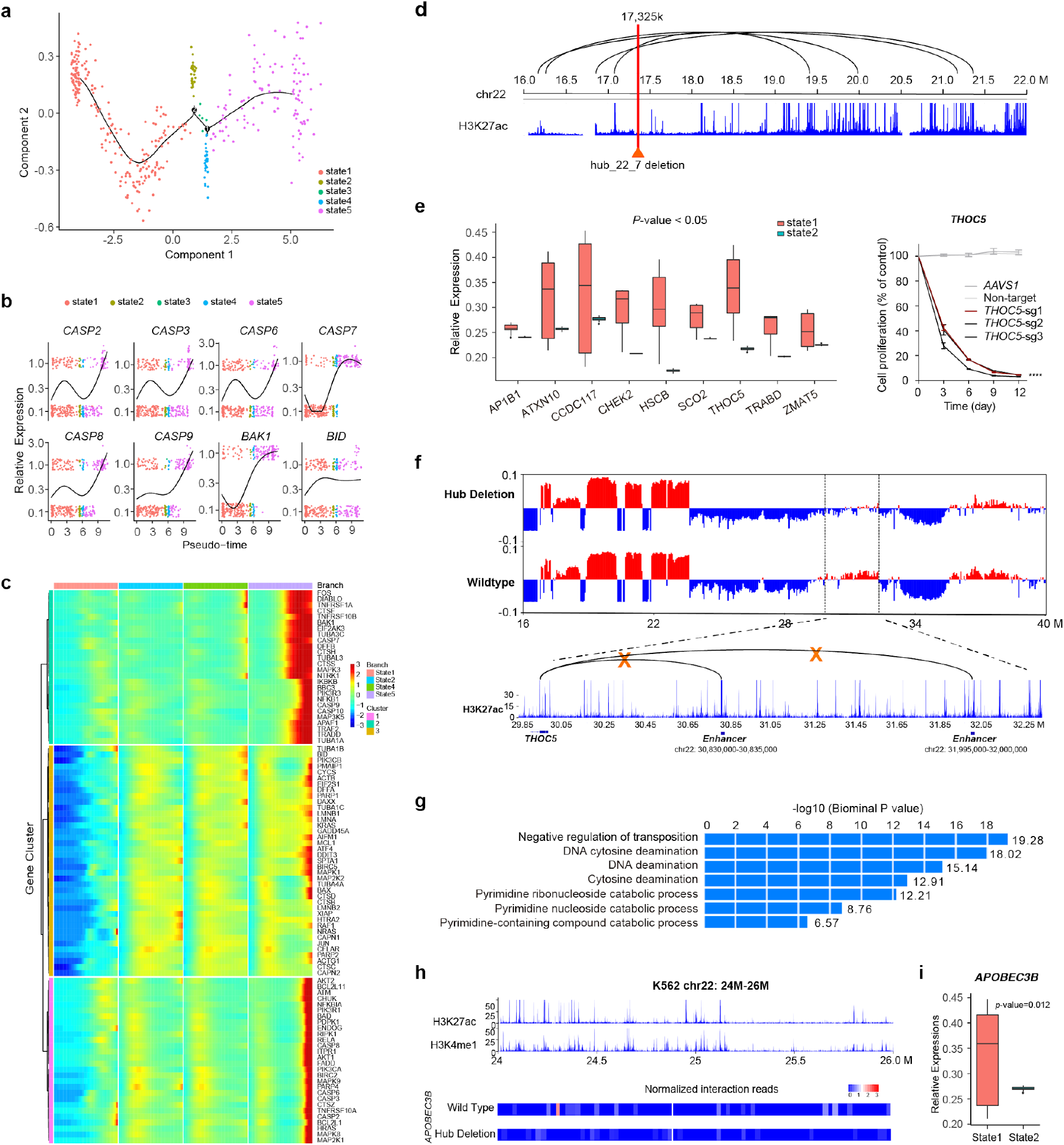
Hub deletion induces global and synergistic change of gene expressions. (**a**) Pseudotime clusters of hub_22_7-deleted and wild type K562 cells based on apoptosis gene expressions. (**b**) Relative expressions of typical apoptosis genes *(CASP2, CASP3, CASP6, CASP7, CASP8, CASP9, BAK1, BID)* under different pseudotime plotted using the Monocle package. Dots in different colors represent different cell states. (**c**) Global analysis of 93 KEGG apoptosis gene expressions in state 1, 2, 4 and 5. Genes were clustered into 3 groups. (**d**) The disruption of enhancers/promoters interactions upon hub_22_7 deletion. (**e**) Essential genes of K562 located in chr22 with significantly down-regulated expression (*p*-value < 0.05) in state2 compared to state1. (**f**) A/B compartment change (50-kb resolution) upon hub deletion. Multiple enhancer-promoter contacts with *THOC5* were disrupted in the compartment changing region (chr22: 29,850,000-32,350,000). (**g**) GO biological process pathways associated with loci whose 3D contacts were disrupted by hub-deletion. (**h**) The 3D contacts between the *APOBEC3B* promoter and enhancers located in chr22: 24,000,000-26,000,000 were significantly decreased in hub-deleted cells. The enhancers were identified using the overlapping peaks of H3K27ac and H3K4me1 in the wild type. (**i**) The relative expression level of *APOBEC3B* from state 1 to 2.

### Deletion of essential hubs can alter gene expressions in distal regions

We noticed that multiple contacts between promoters and enhancers located at the opposite sides of the hub in the linear genome were disrupted upon hub deletion (**Fig. 4d**), indicating that deleting a hub could affect transcriptional regulation. To investigate whether important genes in chr22^26,27^ were affected, we compared the expression profile of state 2 with state1 and found significantly down-regulated genes upon hub deletion, including multiple essential genes in K562 identified from the previous genome-wide screening ^26^, such as *ATXN10, THOC5, CHEK2* and *HSCB* (**Fig. 4e**). Notably, these genes are located distal (12 Mb-34 Mb away in the linear genome) from the deleted hub. We confirmed the essentiality of *THOC5* in K562 through CRISPRi-based gene knockdown (**Fig. 4e**). Furthermore, the high resolution Hi-C data indicated its promoter’s interaction with two enhancers were disrupted upon hub deletion (**Fig. 4f**). Interestingly, they are located in an A (active) to B (inactive) flip region (chr22: 29 Mb-32 Mb, **Fig. 4f**), consistent with *THOC5* repression. These observations suggested that the chromatin structure alteration induced by hub deletion could affect expression of distal genes including those essential for cell viability.

### Hubs can be new non-coding therapeutic targets

Because deleting one essential hub can impact many genes, it suggests a strategy of developing new “one-drug-multiple-targets” therapeutics to synergize different pathways. Namely, the disease-specific non-coding regions, such as hubs that are only essential in cancer cells, could be potential therapeutic targets. In our screening, we did identify hubs specifically essential for K562 (**Fig. 2e-f** and **Supplementary Fig. 4**). As K562 is a leukemia cancer cell line, the K562-specific hubs could be potential therapeutic targets. For example, hub_22_7 deletion resulted in about 80% decrease of cell proliferation rate in K562, but nearly no significant effects in other analyzed cell lines (**Fig. 2f** and **Supplementary Fig. 4a**). As shown above, deletion of this hub caused the down-regulation of many essential genes and the activation of apoptosis pathways. Therefore, this collective effect of killing cancer cells is more potent than targeting each individual pathway, and would make it harder for cancer cells to develop drug resistance.

Furthermore, hub deletion also affects genes specifically expressed in K562 despite they are not essential for cell viability. For example, K562-specifically highly expressed *TOP3B* (**Supplementary Fig. 8**), which plays important roles in the maintenance of gene stabilities and chromosome bridging ^27,28^, was down-regulated upon hub deletion due to the disruption of its promoter-enhancer interactions. By examining the ENCODE data in 23 cell lines/tissues, we found that the enhancers in chr22: 17,125,000-17,130,000 were only marked by H3K27ac in K562 and another leukemia cell line Dnd41 (**Supplementary Fig. 8**). The low expression of *TOP3B* in Dnd41^20^ (**Supplementary Fig. 8**) suggests that these enhancers may only regulate *TOP3B* in K562. Therefore, deleting this hub can specifically down-regulate the *TOP3B* in K562.

We also used GREAT ^29^ to search for pathways enriched (Binomial FDR Q-value ≤ 1E-5) in the loci whose Hi-C contacts (*p*-value ≤ e-20) were significantly reduced upon hub deletion in chr22 (**Fig. 4g**). Notably, the *APOBEC3* family genes stood out and in particular, *APOBEC3B* was significantly down regulated from state 1 to 2 (**Fig. 4i**). This is likely due to the reduced interaction between the *APOBEC3B* promoter and its enhancers upon hub deletion (**Fig. 4h**). *APOBEC3* enzymes were reported as therapeutic targets for cancer treatment ^30,31^ and their aberrant expressions (e.g. higher expression of *APOBEC3B)* could cause cancerous mutagenesis leading to drug resistance or metastasis ^32–34^ Although *APOBEC3B* is not essential for K562 viability, its down-regulation could effectively reduce mutation rate, which is crucial for developing potent therapy. Taken together, deleting one hub is possible to synergize multiple pathways for killing cancer cells and simultaneously reducing the cancer capability to mutate. This example suggests that identification and deletion of cancer-specific hubs could open a new avenue of developing new potent therapeutics.

## DISCUSSION

In this study, we have shown that the 3D structural importance of hubs in the human genome and the essential hubs are not necessarily marked with any epigenomic signal. Deletion of the hubs could alter the expression of genes located very distal in the genome. There could be different ways that hub deletion changes the chromatin structure, such as disrupting the chromatin packing, changing the chromatin loops and promoter-enhancer interactions. Such a change can be propagated from the nearby loci to distal regions. Note that structural importance is not limited to hubs and we also found deleting enhancers could also impact broad chromatin organization despite being less global compared to hub deletion. The non-coding disease-associated GVs can occur in the hub regions to form or disrupt hubs in normal cells, and facilitate the switch to the disease cell state. Our analysis provides a new aspect to understand the mechanisms of non-coding GVs that do not overlap with any functional elements but are tightly associated with diseases. Furthermore, the identified cancer-specific hubs could be potential new therapeutic targets, whose deletion can affect many genes located distal from each other in the genome. Targeting these non-coding loci could leverage the synergistic effects of multiple mechanisms to develop potent therapeutics.

## METHODS

Methods, including statements of data availability and any associated accession codes and references, are available in the online version of the paper.

## ACKNOWLEDGEMENTS

We acknowledge the staff of UC San Diego IGM Genomics Center for sequencing service and UC San Diego Human Embryonic Stem Cell Core Facility for cell sorting service. We acknowledge Ms. Jia Xu (UC San Diego) for the assistance in preparing a single cell RNA-seq library. We acknowledge the staff of the BIOPIC High-throughput Sequencing Center (Peking University) for their assistance in next-generation sequencing analysis, the National Center for Protein Sciences (Beijing) at Peking University for assistance with Fluorescence-activated cell analysis and sorting, and Dr. H. L., Ms. L. D. for technical help. We acknowledge Dr. Ying Yu (Peking University) for the assistance in preparing the NGS library. This project was partially supported by CIRM (RB5-07012) and NIH (R01HG009626) (to Wei Wang), and was supported by funds from the National Science Foundation of China (NSFC31930016), Beijing Municipal Science & Technology Commission (Z181100001318009), the Beijing Advanced Innovation Center for Genomics at Peking University, and the Peking-Tsinghua Center for Life Sciences (to Wensheng Wei).

## AUTHOR CONTRIBUTIONS

W.Wang and W.Wei conceived and supervised the project. W.Wang, W.Wei, B.D. and Y.L. designed the experiments. B.D and L.Z. constructed network analysis, identified and characterized hub regions. Y.G. designed the pgRNA library for hub screening. Y.L. and P.X. performed the pgRNA library construction and screening. Y.L. performed the experiments including individual validation of candidate hubs, differentially expressed genes (DEGs), the whole genome sequencing and bulk RNA-seq with the help of P.X. and Q.P.. Z.L. performed the bioinformatics analysis of the screening data and designed the pgRNAs used for individual validation. P.W and Z.C. performed the Hi-C experiments on hub deletion cell lines. P.W. and Y.Z. performed single cell RNA-seq on hub-deletion cell lines. L.Z. performed the bioinformatics analysis of the single cell RNA-seq and Hi-C data. B.D.,Y.L., L.Z., W.Wei and W.Wang wrote the manuscript with contributions from all other authors.

## ONLINE METHODS

### Network construction and hub identification

#### Evaluating significance of Hi-C interaction pairs

We collected the raw reads, scale factors for vanilla coverage (VC) normalization and the expected normalized reads for interaction pairs from the Hi-C experiments provided by Rao et al. (GSE63525) ^14^ The raw read, *R_ij_*, between fragment *F_i_* and *F_i_* was first divided by both of sequence distance between *F_i_* and *F_j_*, and obtained the expected normalized reads for the scale factors *S_F_i__*. and *S_F_j__*. for VC normalization, 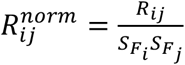. Then we calculated the distance 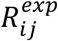. Finally, the significance of the interaction between *F_i_* and *F_j_* was evaluated using the p-value of the normalized read 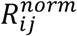 calculated based on Poisson distribution ^4^ with an expectation equal to 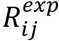.

#### Consideration of translocation in K562

When processing the Hi-C reads in K562, we took the reciprocal translocation between chr9 (9q34) and chr22 (22q11) into account. We mapped reads to chr9 and chr22 in the reference genome and then translocated them. Next, the reads were normalized with the scale factors for each fragment provided by the Rao et al. study^14^. P-value was then calculated in the same way as described above. The translocated chr9 is ~10Mbp longer than the reference chr9. As the expected reads and the genomic distance follow a power-law^4^, we fitted a linear model between logarithm of expected reads and logarithm of genomic distance, and estimated the expected reads for the longer genomic distance.

#### Characterizing hubs

Peaks of six histone modifications (H3K4me1, H3K4me3, H3K27ac, H3K36me3, H3K27me3, and H3K9me3), DNaseI-seq and ChIP-seq for transcription factors (TFs) were counted in the hub regions. The peaks of DNaseI-seq data and ChIP-seq data for TFs were downloaded from the website of ENCODE2 (http://genome.ucsc.edu/encode) when this analysis was done. DNaseI-seq data included 125 cell types and ChIP-seq data included 49, 98, 77 TFs in H1 hESC, GM12878, and K562 cells, respectively. For the peaks of the six core histone modifications, we first collected the data from the NIH Roadmap Epigenomics Project, and then called the peaks following the procedure in ref ^35^: peaks for H3K27ac, H3K4me1, and H3K4me3 were identified using Homer program “findPeaks” with the style “histone”^36^, and peaks within 1-kb were merged into a single peak; peaks for other marks (H3K9me3, H3K36me3 and H3K27me3) were identified by the Homer program “findPeaks” with the parameter options as “-region -size 1000 -minDist 2500”.

For each epigenetic marker, we overlapped their peaks with each hub/non-hub region and counted the number of cell types with overlapping peaks. Distributions of the enrichment were compared between hubs and non-hubs, and p-value was calculated using *Wilcoxon* test.

#### Cell line specificity of the node degree distribution

For 5 kb-resolution Hi-C data in the 5 cell lines, we used correlation-based method to evaluate cell type specificity: 1) the degree of each node was represented as a vector containing the degree z-score values calculated in the 5 cell lines that had both genotype variation and Hi-C data (GM12878, HMEC, HUVEC, IMR90 and K562). 2) For cell type specificities, there are 2^5^=32 possible vectors, including 2 with no cell line specificity (0,0,0,0,0), (1,1,1,1,1), 5 specific to one cell line (1,0,0,0,0), (0,1,0,0,0)…(0,0,0,0,1), 10 specific to two cell lines (1,1,0,0,0), (1,0,1,0,0)…(0,0,0,1,1), 10 specific to three cell lines (1,1,1,0,0), (1,0,1,1,0).(0,0,1,1,1) and 5 specific to four cell lines (1,1,1,1,0), (1,0,1,1,1).(0,1,1,1,1). 3) For each node, we calculated the Pearson correlation between the degree vector and these cell line specificity vectors. If the best correlation coefficient is larger than a threshold of 0.9, we assigned the node with the corresponding cell line specificity.

For 20 kb-resolution Hi-C data in 12 normal cell lines and 2 cancer cell lines, we used distribution-based method to evaluate cell type specificities: 1) the degree of each node was represented as a vector containing the degree z-score values calculated in all cell lines that had both genotype variation and Hi-C data. 2) For each node, we assumed that the normalized degrees obey Gaussian distribution across normal cell lines, and we calculated mean and standard deviation. 3) Based on the mean and standard deviation for each node, we calculated Z-score for each cell line, i.e. the cell line specificity Z-score. The node was considered as cell line specific if the absolute value of cell line specificity Z-score was larger than 1.

### Hub screening and validation

#### Cell culture

K562, H1975 and NAMALWA cells were cultured in RPMI 1640 medium (Gibco). 293T, HeLa, A549 and Huh7.5.1 cells were cultured in Dulbecco’s modified Eagle’s medium (DMEM, Gibco). All cells were supplemented with 10% fetal bovine serum (FBS, Biological Industries) with 1% penicillin/streptomycin, cultured with 5% CO2 in 37°C.

#### Design and construction of the CRISPR-Cas9 pgRNA library

In order to validate the importance of hub regions, we sorted hub regions with PLT, and selected top 700 all-cell-line hubs and top 300 K562-specific hubs. Among them, 960 hubs were suiTable for designing pgRNAs for CRISPR-Cas9 screening. For each hub, up to 20 pgRNAs were designed to target 1-kb upstream and 1-kb downstream regions flanking the two boundaries of the 5-kb segment, respectively. To ensure the cleavage accuracy and efficacy, we required sgRNAs in each pair contain at least 2 mismatches to any other loci in the human genome and their GC contents between 0.2 and 0.8. For all the possible pgRNAs obtained from the selected sgRNAs, we removed those that may delete any promoter or exon of protein-coding genes and we made sure that the cut site of each sgRNA is at least 30 bp away from the exon-intron boundary of coding genes. We also designed 473 pgRNAs deleting promoter region and first exon of 29 ribosomal genes as the positive controls, and 100 pgRNAs targeting *AAVS1* locus, 100 non-targeting pgRNAs as the negative controls, which were obtained from our previous library^8^. As a result, the hub deletion library contained 17,476 pairs of gRNAs targeting 960 hub loci. The 128 nt oligonucleotides containing pgRNA coding sequences were designed, synthesized (Agilent Technologies, Inc.) and cloned into the lentiviral expression vector following the two-step cloning method as previously described^8^, with a minimum representation of 150 transformed colonies per pgRNA in each cloning step.

#### CRISPR-Cas9 pgRNA library screening

K562 cells stably expressing Cas9 were infected with pgRNA library lentiviruses at an MOI of < 0.3 (1000-1500x coverage of the library), and two replicates were arranged. 72 hours after infection, EGFP^+^ cells were selected by FACS (Day-0). For each replicate, the harvested cells were divided into Day-0 control group and experimental group which was further maintained at a minimum coverage of 1500x for 30 days. Then cells of each group with 1500x library coverage were respectively subjected to genomic DNA extraction, PCR amplification of sgRNA-coding sequence and high-throughput sequencing analysis (Illumina HiSeq2500 and HiSeq X Ten platform) as previously described ^8^.

#### Identification of functional hubs

Sequencing reads were mapped to the pgRNA library and further normalized to reads per million (RPM) for each barcode-gRNA. After calculating the quantile of pgRNA counts from two replicates, we removed noisy pgRNAs if a pgRNA’s quantile difference of two replicates was in either 3% tail of the distribution. Then, log_2_ (fold change) (log_2_FC) between experimental and control groups were calculated for each pgRNA, and 100 negative control genes were generated by randomly sampling 20 *AAVS1-targeting* pgRNAs with replacement. Two scores for each set of hub were calculated: 1) the mean log_2_FC of all pgRNA in the set, denoted by *FC_hub_*; 2) −log_10_P_value_ of one-side Mann Whitney U test of all pgRNAs in the set compared with pgRNAs targeting *AAVS1* locus, denoted by *P_hub_*. The background distribution of these two scores was represented by the mean (*μ_FC_* and *μ_p_*) and standard deviation (*σ_FC_* and *σ_p_*) of all negative control genes. Then, the essentiality of hubs was evaluated by the following function:

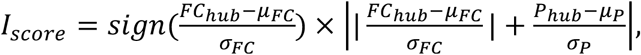

and hubs with lowest *I_score_* (≤ −1) were identified as essential hubs.

To further avoid the potential issue of cell toxicity generated from multiple cleavages by some pgRNAs, we retreived the GuideScan specificity score to evaluate each sgRNA ^19^. By calculating the harmonic mean of the two sgRNAs for each pgRNA, a specificity score was generated for each pgRNA. We only kept the identified essential hubs if their targeting pgRNAs had specificity score is > 0.1 and log_2_ (fold change) is < −1. Furthermore, to avoid the copy number effect on drop-out screening, the copy number of each hub locus in K562 cell line was analyzed based on the ENCODE consortium copy number data (https://www.encodeproject.org/files/ENCFF486MJU/). After further filtering hub loci with copy number amplification, the remaining hits were regarded as essential hubs.

#### Individual validation of essential hubs by cell proliferation assay

For each candidate hub locus, two pgRNAs were used for the individual validations and they were either newly designed or selected from the library showing consistent depletion in replicates. To ensure high targeting specificity of all the selected pgRNAs, we required that their specificity scores are all larger than 0.15, and the score of at least one pgRNA for each hub is larger than 0.2. For the newly-designed pgRNA, we further required that they do not include ≥ 4-bp homopolymer stretch and their GC contents are between 0.4 and 0.7. We also changed the deletion regions which include each sgRNA targeting −1-kb to +0.5-kb flanking the two boundaries of the 5kbp hub loci (− and + refer to outer and inner hub direction, respectively). Other rules were the same as those used for pgRNA design of the library screening.

All the pgRNAs targeting each hub to be validated were individually cloned into a lentiviral expression vector containing an EGFP selection marker. After virus packaging, the pgRNA lentiviruses were respectively transduced into K562 cells at an MOI of < 1. The cell proliferation assay was performed as previously described ^8,10^. The experiments lasted for 15 days after the first FACS analysis and at least 100,000 cells were analyzed.

#### Whole genome sequencing (WGS) to evaluate off-target effect

K562 cells were lentivirally transduced with the pgRNA hub_22_7-pg2. The EGFP positive cells were collected by FACS sorting at Day 8 post pgRNA infection at an MOI of < 1, and the sorted cells were subjected to genomic DNA extraction. The whole genome sequencing (WGS) library was prepared following the manufacturer’s instruction and was sequenced using the Illumina HiSeq 4000 platform. Using the WGS data, we evaluated the deletion efficiency in the targeted locus and off-target effects.

We downloaded the K562 (wild type) WGS data from ENCODE with accession code ENCFF313MGL, ENCFF004THU, ENCFF506TKC and ENCFF066GQD, and then evaluated the potential off-target effects following the published procedures ^37,38^. We first generated the putative off-target sites for hub_22_7 in the hg19 genome using Cas-OFFinder ^37^. We called base mismatch type with at most 4 mismatches without considering any bulge (mismatch ≤ 4, bulge = 0). We also called bulge mismatch type with at most 2 mismatches with at maximum 2 bulges (mismatch ≤ 2, bulge ≤ 2). In total we examined 455 potential off-target loci. In order to detect the candidate mutations and indels in the hub deleted cells, variant calling was performed as described on GATK Best Practices (https://gatk.broadinstitute.org/hc/en-us). Briefly, reads were aligned to the human reference genome (hg19) using BWA-0.7.17. Duplicated reads were then removed using GATK4 MarkDuplicatesSpark(https://gatk.broadinstitute.org/hc/en-us/articles/360037224932-MarkDuplicatesSpark).The reads were then processed via base quality score recalibration using GATK4. Germline mutations (compared to the hg19 reference genome) were called in both wild type and hub-deleted cells by GTAK HaplotypeCaller (version 4.1.4.1) with default parameters. SNVs and Indels called by GATK4 Mutect2 (version 4.1.4.1) with default parameters were used to assess off-target deletion.

We further confirmed no off-target effect using a different analysis software, BCFTOOLS suite (version 1.9, http://www.htslib.org/doc/bcftools.html), to re-examine the SNP and indel site from the WGS data. The mapped BAM file of K562 cells was piped into bcftools mpileup and bcftools call with default parameters. The called raw VCF file was filtered by bcftools filter with “%QUAL<30 || DP < 30” marked as low-quality variants. Homozygous variants were also removed from the raw VCF file with parameter “GT=1/1”. Gold standard indels VCF of Mills and 1000G was downloaded from GATK Resource Bundle (https://gatk.broadinstitute.org/hc/en-us/articles/360036212652-Resource-Bundle). The gold standard indels were also removed from the VCF file using bcftools isec with parameter “-n −1 -c all”. There were no putative off-target sites found in the 13809 indels obtained using bedtools intersect (https://bedtools.readthedocs.io).

### Hi-C library preparation and data analysis

#### Hi-C library preparation

The pgRNA Hub_22_7-pg2 was delivered into K562 cells through lentiviral infection at an MOI of < 1. The EGFP positive cells were collected by FACS sorting at Day 9 post infection and the sorted cells were allowed to recover in normal cell culture condition for 2 hr before proceeding to Hi-C library. One million cells were used for each Hi-C library preparation using Arima-HiC kit (Arima Genomics, San Diego) following the manufacturer’s instruction. Hi-C libraries were sequenced using the Illumina NovaSeq platform.

#### Hi-C data processing

The Hi-C raw fastq data were processed by Juicer pipeline^39^ with default parameters. Hi-C reads were aligned to hg19 (GRCh37) and the reads with MAPQ < 30 were further trimmed (**Supplementary Table 14**). The output bam files were transformed into 5kb, 10-kb, 25-kb, 50-kb, 100-kb and 1-Mb resolution contact matrix. The contact matrix was then normalized by the vanilla coverage (VC) method^14^. The significance level of a given interaction pair was calculated from Poisson Distribution fitting between the measured interaction reads and the expected reads by VC normalization. Juicebox (https://www.aidenlab.org/juicebox/) and HiCExplorer ^40,41^ was utilized to visualize the processed Hi-C data.

#### Loop calling

In both wild type K562 and hub_22_7-deleted K562 cells, the VC normalized Hi-C contact reads were processed by HiCCUPS at 25kb resolution for calling loops.

(https://github.com/aidenlab/juicer/wiki/HiCCUPS). Loops were called using a FDR-cutoff (bottom_left, donut, horizontal, vertical) of 10^−5^.

#### A/B compartment analysis

The A/B compartment analysis was conducted using 50-kb bins. The eigenvectors for each chromosome in both K562 wildtype and hub deleted cells were extracted from the VC normalized Hi-C counts processed by the Juicer pipeline with default parameters^39^. The Pol II ChIP-seq data in K562 was downloaded from ENCODE ^42^. The correlation between the first eigenvector of each chromosome and the Pol II peaks was calculated, based on which we determined the A and B compartment ^43^.

#### Effective diameter comparison

We calculated the effective diameter deviation for each chromosome both before and after hub deletion, and found that the deviation followed Gaussian distribution by Shapiro-Wilk normality test (*p*-value=0.27 so that the null hypothesis of being normal distribution was accepted). Then we calculated the *p*-value for the deviation of each chromosome based on Gaussian distribution, and identified the significant changed chromosome with *p*-value < 0.05.

#### Modularity comparison

We collected the modularity scores of each chromosome in the seven wild type cell lines (GM12878, K562, HUVEC, IMR90, NHEK, KBM7 and HMEC), and found that the modularity score for each chromosome followed Gaussian distributions (all the p-value ≥ 0.01 to accept the null hypothesis of being a Gaussian distribution in Shapiro-Wilk normality test). Then for each chromosome in hub-deleted K562, we calculated the *p*-value of its modularity score based on chromosome-specific modularity distribution and identified significant changed chromosomes with p-value < 0.05.

### Bulk RNA-seq and data analysis

#### Bulk RNA-seq library preparation

The pgRNA *AAVS1* -pg1 targeting *AAVS1* locus was delivered into K562 cells at an MOI of < 1. 2×10^6^ EGFP+ K562 cells were sorted by FACS 8 days post infection. The total RNA was extracted using RNeasy Mini Kit (QIAGEN 79254), and three replicates were arranged. The RNA-seq libraries were further prepared following the NEBNext PolyA mRNA Magnetic Isolation Module (NEB E7490S), NEBNext RNA First Strand Synthesis

Module (NEB E7525S), NEBNext mRNA Second Strand Synthesis Module (NEB E6111S) and NEBNext Ultra DNA Library Prep Kit for Illumina (NEB E7370L). All samples were subjected to NGS analysis using the Illumina HiSeq 4000 platform.

#### Bulk RNA-seq data processing

In the bulk RNA-seq library, the sequencing reads with Phred score >=30 were aligned to the human reference genome (GRCh37/hg19) using HISAT2 (2.0.4)^44,45^ and assembled and quantified by StringTie (1.3.5) ^44,46^. The gene read counts for each sample were further normalized by transcripts per kilobase million (TPM).

### Single cell RNA-seq and data analysis

#### Single cell library preparation

K562 cells infected with Hub_22_7-pg2 were FACS-sorted 8 days after lentivirus transduction for single-cell library preparation. The single-cell library was prepared with established protocol described previously^22^. Briefly, polyA+ RNA was reverse transcribed through tailed oligo-dT priming directly in whole cell lysate (single droplet) using Moloney Murine Leukemia Virus Reverse Transcriptase (MMLV RT) and temperature switch oligos. The resulting full-length cDNA contains the complete 5’ end of the mRNA, as well as an anchor sequence that serves as a universal priming site for second strand synthesis. The cDNA was pre-amplified using 15 cycles with Kapa HiFi Hotstart Readymix. We used the Nextera DNA sample preparation kit to generate the single cell libraries. The amplified cDNA was tagmented at 55°C for 5 min in 20 μl reaction with 0.25 μl of transposase and 5 μl Nextera reaction buffer. 35 μl of PB was added to the tagmentation reaction mix to strip the transposase off the DNA, and the tagmented DNA was purified with 20 μl of Ampure beads (sample to beads ratio of 1:0.6). Purified DNA was then amplified by 12 cycles of standard Nextera PCR. The prepared libraries were sequenced on Illumina HiSeq4000.

#### Single cell RNA-seq processing

The fastq files were first mapped to the human reference genome (GRCh37/hg19) using Picard (2.17.0)^47^ and STAR(2.5.3a)^48^. We used the Drop-seq processing pipeline developed by McCarroll lab ^22^ to remove low quality reads (lower than Q10) and PCR duplicates (identified by cell-barcode and molecular-barcode). The cells were ordered descendingly by the read count. Reads from all the cells were pooled together to form a cumulative distribution. Cells with the most reads before the inflection point “knee” of the cumulative distribution were kept for the following analysis (**Supplementary Table 15**).

We calculated a p-value for each gene to assess whether the change was significant or not. Each cell was first normalized by counts per million (CPM). We calculated *E_i_*, which is the sum of CPMs for a given gene across all the cells, and *E_total_,* which is the sum of *E_i_* for all the genes. We then computed *P_i_=E_i_* / *E_total_*. In a given cell *j*, the normalized gene expressions of all genes were assumed to follow a binomial distribution *G_ij_~B (N_j_,P_i_*) independently identically, where *G_ij_* is the expected reads of the gene *i* in cell *j*, *N_j_* is the total reads for cell *j*. We calculated a p-value to evaluate how significantly each gene expression in each cell deviated from the expected value based on the binomial distribution, which indicates its differential expression across cells. We also calculated *p*-value for genes in the negative control *(ΔAAVS1)* and wild type bulk RNA-seq data in the same way.

#### Single-cell trajectory branching and pseudotime analysis

Because hub deletion affected cell proliferation, we focused on analyzing the apoptosis genes annotated in the KEGG database ^49^. Considering the noise in the single cell RNA-seq data, we selected the apoptosis genes that showed differential expression in at least 10% ~15% cells (*p*-value<0.05). As a result, 93 apoptosis genes were identified in K562 cells with the essential hub chr22: 17,325,000-17,330,000 deleted. All the single cell and bulk data were clustered with trajectory branching and pseudotime analysis using the Monocle R package ^23,24^ Monocle^23,24^ assigned each cell a pseudotime value and a “State” on the basis of the segment of the trajectory according to the PQ-tree algorithm. Cells with the same “State” were clustered together ^24^ and then relative gene expressions in each cluster were computed.

#### Differentially expressed genes identified from pseudotime analysis

To identify differentially expressed genes (DEGs) between state 1 and state 2 defined in the pseudotime analysis, a *Wilcoxon* Rank-Sum Test was applied to identify genes significantly down regulated in state 2 compared to state 1.

#### Investigation of the essentialities of differentially expressed genes from single cell RNA-seq data

Among the differentially expressed genes (DEGs) in chr22 upon hub_22_7 (chr22: 17,325,000-17,330,000) deletion, which were significantly decreased from state 1 to state 2, a topranked DEG *THOC5* was selected to analyze its importance on cell growth and proliferation in K562 cells. Three sgRNAs were designed to knock down its expression through CRISPRi strategy, which were selected from the hCRISPRi-v2 library ^26^. These sgRNAs were also individually cloned into the lentiviral expression vector with an EGFP marker, and then respectively transduced into K562 cells stably expressing dCas9-KRAB protein at an MOI of <1. The cell proliferation assay was performed as previously described ^8,10^. The first time point of FACS analysis was at 6 days post lentiviral infection, and the experiment lasted for 12 days.

## REFERENCES

1. Dixon, J. R. et al. Topological domains in mammalian genomes identified by analysis of chromatin interactions. Nature 485, 376–380 (2012).

2. Nora, E. P. et al. Spatial partitioning of the regulatory landscape of the X-inactivation centre. Nature 485, 381–385 (2012).

3. Lupiáñez, D. G. et al. Disruptions of topological chromatin domains cause pathogenic rewiring of gene-enhancer interactions. Cell 161, 1012–1025 (2015).

4. Lieberman-Aiden, E. et al. Comprehensive mapping of long-range interactions reveals folding principles of the human genome. Science 326, 289–293 (2009).

5. Dekker, J., Marti-Renom, M. A. & Mirny, L. A. Exploring the three-dimensional organization of genomes: interpreting chromatin interaction data. Nat. Rev. Genet. 14, 390–403 (2013).

6. Nagano, T. et al. Single-cell Hi-C reveals cell-to-cell variability in chromosome structure. Nature 502, 59–64 (2013).

7. Rose, G. D., Fleming, P. J., Banavar, J. R. & Maritan, A. A backbone-based theory of protein folding. Proc. Natl. Acad. Sci. U. S. A. 103, 16623–16633 (2006).

8. Zhu, S. et al. Genome-scale deletion screening of human long non-coding RNAs using a paired-guide RNA CRISPR-Cas9 library. Nat. Biotechnol. 34, 1279–1286 (2016).

9. Liu, S. J. et al. CRISPRi-based genome-scale identification of functional long noncoding RNA loci in human cells. Science 355, (2017).

10. Liu, Y. et al. Genome-wide screening for functional long noncoding RNAs in human cells by Cas9 targeting of splice sites. Nat. Biotechnol. (2018) doi:10.1038/nbt.4283.

11. Fulco, C. P. et al. Systematic mapping of functional enhancer-promoter connections with CRISPR interference. Science 354, 769–773 (2016).

12. Simeonov, D. R. et al. Discovery of stimulation-responsive immune enhancers with CRISPR activation. Nature 549, 111–115 (2017).

13. Diao, Y. et al. A tiling-deletion-based genetic screen for cis-regulatory element identification in mammalian cells. Nat. Methods 14, 629–635 (2017).

14. Rao, S. S. P. et al. A 3D map of the human genome at kilobase resolution reveals principles of chromatin looping. Cell 159, 1665–1680 (2014).

15. Munoz, D. M. et al. CRISPR Screens Provide a Comprehensive Assessment of Cancer Vulnerabilities but Generate False-Positive Hits for Highly Amplified Genomic Regions. Cancer Discovery vol. 6 900–913 (2016).

16. Aguirre, A. J. et al. Genomic Copy Number Dictates a Gene-Independent Cell Response to CRISPR/Cas9 Targeting. Cancer Discov. 6, 914–929 (2016).

17. Morgens, D. W. et al. Genome-scale measurement of off-target activity using Cas9 toxicity in high-throughput screens. Nat. Commun. 8, 15178 (2017).

18. Tycko, J. et al. Mitigation of off-target toxicity in CRISPR-Cas9 screens for essential noncoding elements. Nat. Commun. 10, 4063 (2019).

19. Perez, A. R. et al. GuideScan software for improved single and paired CRISPR guide RNA design. Nat. Biotechnol. 35, 347–349 (2017).

20. The ENCODE Project Consortium. An integrated encyclopedia of DNA elements in the human genome. Nature 489, 57–74 (2012).

21. Bae, S., Park, J. & Kim, J.-S. Cas-OFFinder: a fast and versatile algorithm that searches for potential off-target sites of Cas9 RNA-guided endonucleases. Bioinformatics 30, 1473–1475 (2014).

22. Macosko, E. Z. et al. Highly Parallel Genome-wide Expression Profiling of Individual Cells Using Nanoliter Droplets. Cell 161, 1202–1214 (2015).

23. Trapnell, C. et al. The dynamics and regulators of cell fate decisions are revealed by pseudotemporal ordering of single cells. Nat. Biotechnol. 32, 381–386 (2014).

24. Qiu, X. et al. Single-cell mRNA quantification and differential analysis with Census. Nat. Methods 14, 309–315 (2017).

25. Kanehisa, M. et al. Data, information, knowledge and principle: back to metabolism in KEGG. Nucleic Acids Res. 42, D199–205 (2014).

26. Horlbeck, M. A. et al. Compact and highly active next-generation libraries for CRISPR-mediated gene repression and activation. Elife 5, (2016).

27. Wang, T. et al. Identification and characterization of essential genes in the human genome. Science 350, 1096–1101 (2015).

28. Zhang, T. et al. Loss of TOP3B leads to increased R-loop formation and genome instability. Open Biology vol. 9 190222 (2019).

29. McLean, C. Y. et al. GREAT improves functional interpretation of cis-regulatory regions. Nat. Biotechnol. 28, 495–501 (2010).

30. Venkatesan, S. et al. Perspective: APOBEC mutagenesis in drug resistance and immune escape in HIV and cancer evolution. Ann. Oncol. 29, 563–572 (2018).

31. Olson, M. E., Harris, R. S. & Harki, D. A. APOBEC Enzymes as Targets for Virus and Cancer Therapy. Cell Chem Biol 25, 36–49 (2018).

32. Swanton, C., McGranahan, N., Starrett, G. J. & Harris, R. S. APOBEC Enzymes: Mutagenic Fuel for Cancer Evolution and Heterogeneity. Cancer Discov. 5, 704–712 (2015).

33. Roper, N. et al. APOBEC Mutagenesis and Copy-Number Alterations Are Drivers of Proteogenomic Tumor Evolution and Heterogeneity in Metastatic Thoracic Tumors. Cell Rep. 26, 2651–2666.e6 (2019).

34. Zhang, Y., Delahanty, R., Guo, X., Zheng, W. & Long, J. Integrative genomic analysis reveals functional diversification of APOBEC gene family in breast cancer. Hum. Genomics 9, 34 (2015).

35. Whitaker, J. W., Chen, Z. & Wang, W. Predicting the human epigenome from DNA motifs. Nat. Methods 12, 265–72, 7 p following 272 (2015).

36. Heinz, S. et al. Simple Combinations of Lineage-Determining Transcription Factors Prime cis-Regulatory Elements Required for Macrophage and B Cell Identities. Mol. Cell 38, 576–589 (2010).

37. Kim, D. et al. Digenome-seq: genome-wide profiling of CRISPR-Cas9 off-target effects in human cells. Nat. Methods 12, 237–43, 1 p following 243 (2015).

38. Smith, C. et al. Whole-genome sequencing analysis reveals high specificity of CRISPR/Cas9 and TALEN-based genome editing in human iPSCs. Cell Stem Cell 15, 12–13 (2014).

39. Durand, N. C. et al. Juicer Provides a One-Click System for Analyzing Loop-Resolution Hi-C Experiments. Cell Syst 3, 95–98 (2016).

40. Ramírez, F. et al. High-resolution TADs reveal DNA sequences underlying genome organization in flies. Nat. Commun. 9, 189 (2018).

41. Wolff, J. et al. Galaxy HiCExplorer: a web server for reproducible Hi-C data analysis, quality control and visualization. Nucleic Acids Res. 46, W11–W16 (2018).

42. Yip, K. Y. et al. Classification of human genomic regions based on experimentally determined binding sites of more than 100 transcription-related factors. Genome Biol. 13, R48 (2012).

43. Kalhor, R., Tjong, H., Jayathilaka, N., Alber, F. & Chen, L. Genome architectures revealed by tethered chromosome conformation capture and population-based modeling. Nat. Biotechnol. 30, 90–98 (2011).

44. Pertea, M., Kim, D., Pertea, G. M., Leek, J. T. & Salzberg, S. L. Transcript-level expression analysis of RNA-seq experiments with HISAT, StringTie and Ballgown. Nat. Protoc. 11, 1650–1667 (2016).

45. Kim, D., Langmead, B. & Salzberg, S. L. HISAT: a fast spliced aligner with low memory requirements. Nat. Methods 12, 357–360 (2015).

46. Pertea, M. et al. StringTie enables improved reconstruction of a transcriptome from RNA-seq reads. Nat. Biotechnol. 33, 290–295 (2015).

47. Picard Tools - By Broad Institute. https://broadinstitute.github.io/picard/.

48. Dobin, A. et al. STAR: ultrafast universal RNA-seq aligner. Bioinformatics 29, 15–21 (2013).

49. Kanehisa, M., Furumichi, M., Tanabe, M., Sato, Y. & Morishima, K. KEGG: new perspectives on genomes, pathways, diseases and drugs. Nucleic Acids Res. 45, D353–D361 (2017).

